# Robust trigger wave speed in *Xenopus* cytoplasmic extracts

**DOI:** 10.1101/2023.12.22.573127

**Authors:** Jo-Hsi Huang, Yuping Chen, William Y. C. Huang, Saman Tabatabaee, James E. Ferrell

## Abstract

Self-regenerating trigger waves can spread rapidly through the crowded cytoplasm without diminishing in amplitude or speed, providing consistent, reliable, long-range communication. The macromolecular concentration of the cytoplasm varies in response to physiological and environmental fluctuations, raising the question of how or if trigger waves can robustly operate in the face of such fluctuations. Using *Xenopus* extracts, we found that mitotic and apoptotic trigger wave speeds are remarkably invariant. We derived a model that accounts for this robustness and for the eventual slowing at extremely high and low cytoplasmic concentrations. The model implies that the positive and negative effects of cytoplasmic concentration (increased reactant concentration vs. increased viscosity) are nearly precisely balanced. Accordingly, artificially maintaining a constant cytoplasmic viscosity during dilution abrogates this robustness. The robustness in trigger wave speeds may contribute to the reliability of the extremely rapid embryonic cell cycle.

## INTRODUCTION

Frog eggs are large cells that are particularly well-suited to quantitative biochemical studies. The eggs are about 1.3 mm in diameter and 1 µL in volume, which makes them amenable to single-cell biochemical assays^1^. Moreover, they can be lysed with minimal dilution, and the undiluted cytoplasm can be recovered and studied^2,3^. These egg extracts self-organize into cell- like compartments^4^, and like the cells from which they are derived, they can carry out rapid cell cycles^2,5,6^ and, under adverse conditions, die by apoptosis^7,8^. Indeed, *Xenopus* egg extracts have provided important insights into the regulation of both the cell cycle and apoptosis.

The large size of the frog egg presents a challenge shared by other large cells and tissues: how to coordinate rapid processes like mitotic entry and apoptotic death across such large distances. Early modeling work on the cell cycle suggested that mitosis might spread through the egg via trigger waves of Cdk1 activity^9^. Trigger waves can occur in systems with positive feedback loops, and they spread faster over large distances than diffusion alone would allow^10,11^. Experimental work has shown that mitosis does spread through *Xenopus* cytoplasm via trigger waves^5,12^, at a speed of ∼60 µm min^-1^, and apoptosis does as well, at about half that speed^8^. A growing body of evidence suggests that trigger waves may be a common way of transmitting signals over large distances in biological systems. Action potentials and calcium waves are familiar examples of trigger waves, as are intercellular cAMP waves in swarming *Dictyostelium*^13–15^ and intercellular ERK waves in wounded fish scales^16^ and mouse skin^17^. Recent work suggests that the remarkable regeneration of an amputated planarian depends upon signals transmitted from the wound site via intercellular trigger waves of ERK activation^18^.

The cytoplasm is a crowded, spatially organized mixture of organelles, macromolecules, and small molecules. Protein concentrations in *Xenopus* extracts^19–21^ and mammalian cell lines^22^ are typically on the order of 75 mg mL^-1^, although it is higher in some cells, e.g. erythrocytes. It has been conjectured that the nominal cytoplasmic protein concentration maximizes the speed of important biochemical processes^23^. The extent to which this conjecture holds true for the cells awaits experimental investigations. Conversely, protein concentration is dynamic; it falls by ∼15% when cells enter mitosis^24,25^ and by ∼50% when cells become senescent^26^. Cells change in volume when attaching to substrates of different stiffnesses^27,28^, and recent work indicates that neutrophils swell by ∼15% in response to chemoattractants, and that the swelling facilitates rapid migration^29^. The extent to which changes in volume, and changes in cytoplasmic concentration, impact the biochemistry of living cells is as yet poorly understood.

Here we ask how long-range communication via trigger waves is affected by changes in the concentration of cytoplasmic *Xenopus* egg extracts. We show that both mitotic and apoptotic trigger waves can be generated and propagated over a wide range of cytoplasmic concentrations. The wave speeds are maximal or near maximal at a 1x cytoplasmic concentration, in line with Dill’s conjecture that the nominal 1x concentration maximizes the speeds of critical biochemical processes^21,23^, and in the case of apoptotic trigger waves the speed is almost invariant over concentrations from 0.1x to 2x. We derive a simple general equation for trigger wave speed as a function of cytoplasmic concentration, which shows how balanced opposing effects are responsible for this robustness, and show that the equation satisfactorily accounts for our experimental observations. Finally, we show that disrupting the balance by maintaining a constant viscosity when diluting the extracts makes trigger wave speed highly sensitive to cytoplasmic concentration.

## RESULTS

### Mitotic trigger waves in concentrated and diluted extracts

Mitosis is brought about by a complex, interconnected regulatory system centered on a protein kinase, cyclin B-Cdk1, and two opposing phosphatases, PP1 and PP2A-B55 (Fig. 1a). Several positive feedback and double-negative feedback loops are embedded in this regulatory system; for example, active cyclin B-Cdk1 turns on its activator Cdc25, and cyclin B-Cdk1 and PP2A-B55 antagonize each other via intertwined double-negative feedback loops (Fig. 1a). The net result of these feedback loops is that the system functions as a bistable switch^30,31^, and this bistability is key for the propagation of the mitotic state as a trigger wave^12^.

**Figure 1.**
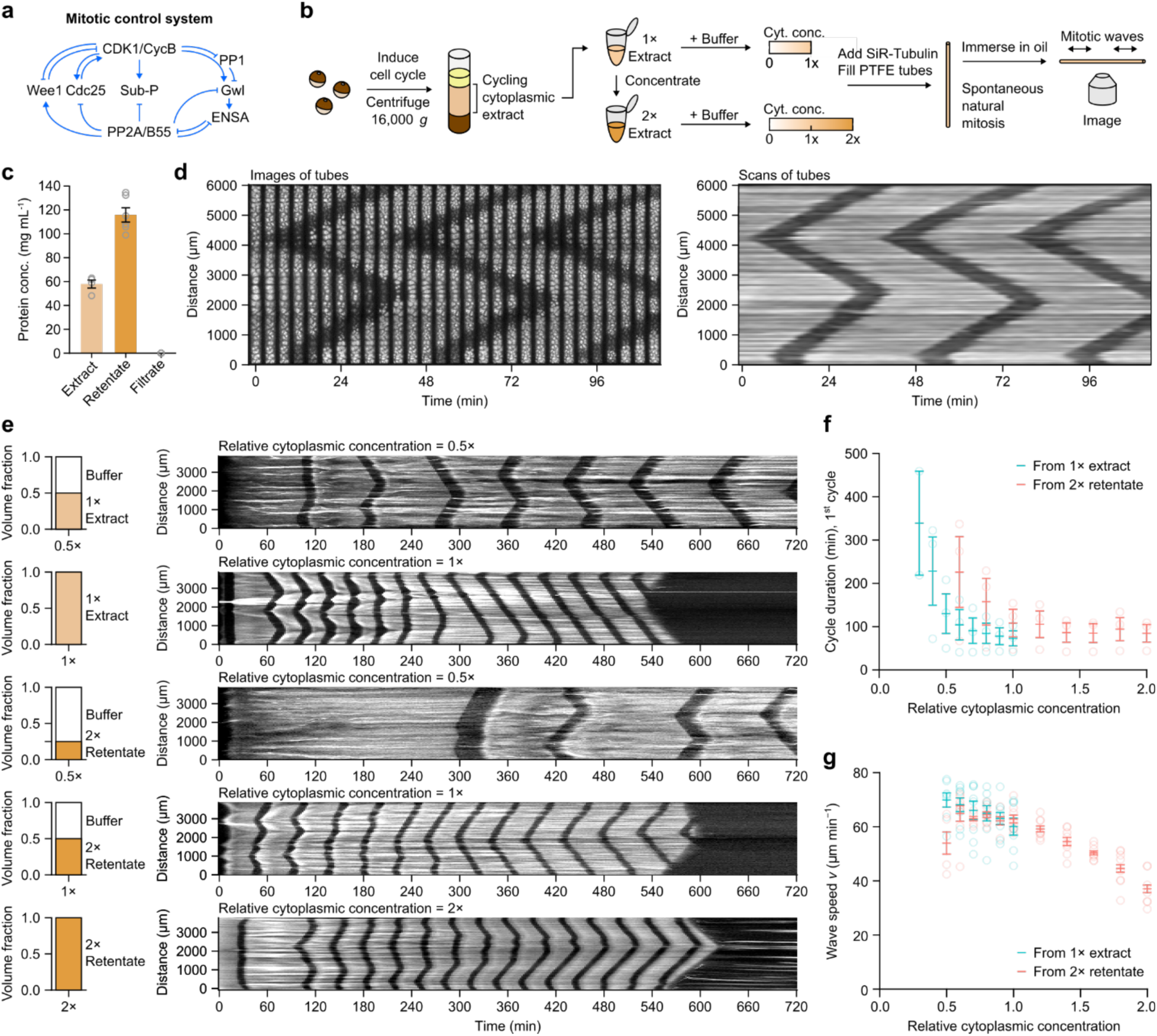
Mitotic trigger wave speed is robust to change in cytoplasmic concentration. **a** Schematic view of the mitotic control network. Note the multiple interconnected positive and double-negative feedback loops. **b** Preparation of cycling *Xenopus* egg extracts and the workflow of monitoring spontaneous mitotic trigger waves in thin PTFE tubes by epifluorescence microscopy. **c** Measurements of protein concentrations in the original and concentrated extract (retentate) as well the filtrate from the ultrafiltration filters. **d** A montage (left) and its corresponding kymograph (right) of a single tube undergoing 3 rounds of mitosis in the course of ∼2 hours. The bright signal is polymerized microtubules stained with SiR-tubulin and the dark bands correspond to the mitotic state in which most microtubules are depolymerized. **e** Representative kymographs (right column) of extracts of different cytoplasmic concentrations prepared from 1x extracts (top 2 rows) or from 2x retentate (bottom 3 rows). Left column shows the volume fractions of buffer (XB buffer without sucrose) and extracts that went into the samples. **f** Duration of the first completely observable cell cycle under the microscope, starting from interphase. Means ± S.E.M. are shown, n = 3 (three independent experiments). **g** Speeds of mitotic trigger waves at different cytoplasmic concentrations. Means ± S.E.M. are shown. Data is compiled from 8 independent experiments.

To see how robust mitotic trigger waves are to changes in the concentration of the cytoplasm, we began by making either a 1x cytoplasmic extract or a concentrated extract on a Microcon spin column. From 4 independent preparations, the 1x cytoplasmic protein concentration was 57.9 ± 3.4 mg mL^-1^ (mean ± S.E.M., n = 4; Fig. 1c), in line with other estimates^19–21^, and not far from the protein concentrations measured for three common mammalian cell lines (∼75 mg mL^-1^)^22^. The concentrated extract was 116 ± 6.0 mg mL^-1^ (mean ± S.E.M., n = 6; Fig. 1c); hereafter we will refer to it as a 2x retentate. The flow-through, which we will refer to as the filtrate, from the spin column had a protein concentration of less than 0.01 mg mL^-1^ (Fig. 1c).

We then diluted the 1x extract or 2x retentate to various extents. In other recent work we used filtrate for the dilutions^21^. Here we have used XB buffer without sucrose rather than filtrate, which allowed us to produce larger volumes of diluted extracts, and we verified that the behaviors of the buffer-diluted and filtrate-diluted extracts were similar (Supplementary Fig. 1).

We added demembranated sperms and SiR-tubulin to the extracts and dilution buffers, made the dilutions, and aspirated extracts into ∼100 µm or ∼300 µm inside-diameter polytetrafluorethylene (PTFE) tubes under gentle vacuum. The tubes were placed under mineral oil and followed by fluorescence video microscopy. Fig. 1d, left panel, shows a typical result. At the first time point shown here, the extract was in interphase with stable microtubules throughout the length of the tube. Within a few minutes, mitosis began near the bottom of the tube and at a locus about 4 mm up the tube. As judged by the depolymerization of the fluorescent interphase microtubules, mitosis spread outward from these two loci in a linear fashion (Fig. 1d). Mitotic exit followed about 12 min after mitotic entrance, and it also spread linearly outward from the same two locations. Fig. 1d, right panel, shows the same data where instead of imaging the whole tube, we recorded SiR-tubulin fluorescence intensity along a line down the middle of the tube and then assembled the data into a kymograph. In either representation, the trigger wave character of mitotic propagation is apparent, and the speed of the mitotic front was 60.2 ± 3.2 µm min^-1^ (mean ± S.E.M., n = 9), similar to previously reported mitotic wave speeds^5,12,32^.

Next we examined how the cell cycle period and the speed of the mitotic waves were affected by changes in cytoplasmic concentration. Fig. 1e shows examples of kymographs from a diluted 1x extract and diluted 2x retentate. The periods of the first cycles and the wave speeds were calculated and are summarized in Figs. 1f and 1g, which include multiple experiments and more dilutions. Several general trends are apparent. First, the wave speeds were similar for 1x extracts and 2x retentates diluted back to 1x, but the periods were different, with the diluted 2x retentates having longer cell cycle periods than the corresponding 1x and diluted 1x extracts. Second, the cell cycle periods tended to be longer in diluted extracts than in concentrated extracts (Fig. 1f). Third, the diluted extracts tended to live longer than the concentrated extracts; a wave of apoptosis, which destroys the microtubule fluorescence, can be seen in the second and in fourth kymographs (Fig. 1e). Fourth, the most concentrated extracts tended to arrest in mitosis with depolymerized microtubules (Supplementary Fig. 2; cf. Fig. 1e, where the extract did not arrest in mitosis). And finally, the speeds of the mitotic waves were relatively invariant, with only the extracts at greater than 1x showing some slowing of the waves. Diluting the extract below 1x slightly increased the wave speed (by ∼10%). For comparison, if the wave speed were determined by a bimolecular reaction, decreasing the extract from 1x to 0.5x might be expected to decrease the wave speed by 75%. Both the cell cycle frequency and mitotic wave speed were at or near their maximal values at 1x cytoplasmic concentration, consistent with Dill’s conjecture^23^.

### Apoptotic trigger waves in concentrated and diluted extracts

Apoptosis is mediated by a complex system of regulators that bring about the activation of caspases 3 and 7, so-called executioner caspases that cleave diverse cellular proteins and bring a halt to the basic processes of life (Fig. 2a). There are several potential positive feedback loops in the apoptotic control system (Fig. 2a), raising the possibility that caspase activation could spread via trigger waves. In many cell types, apoptosis spreads through the cytoplasm in a wave-like manner^33–35^, and in *Xenopus* egg extracts, where it is easy to obtain length scales over which the distinction between diffusive spread and trigger wave spread is unambiguous, it is clear that the fronts of caspase activation represent trigger waves that propagate without slowing down or decreasing in amplitude^8^. The manipulability of extracts allowed us to assess the sensitivity of the apoptotic trigger wave speed in such extracts to cytoplasmic concentration, and to tease out the contributing effects quantitatively.

**Figure 2.**
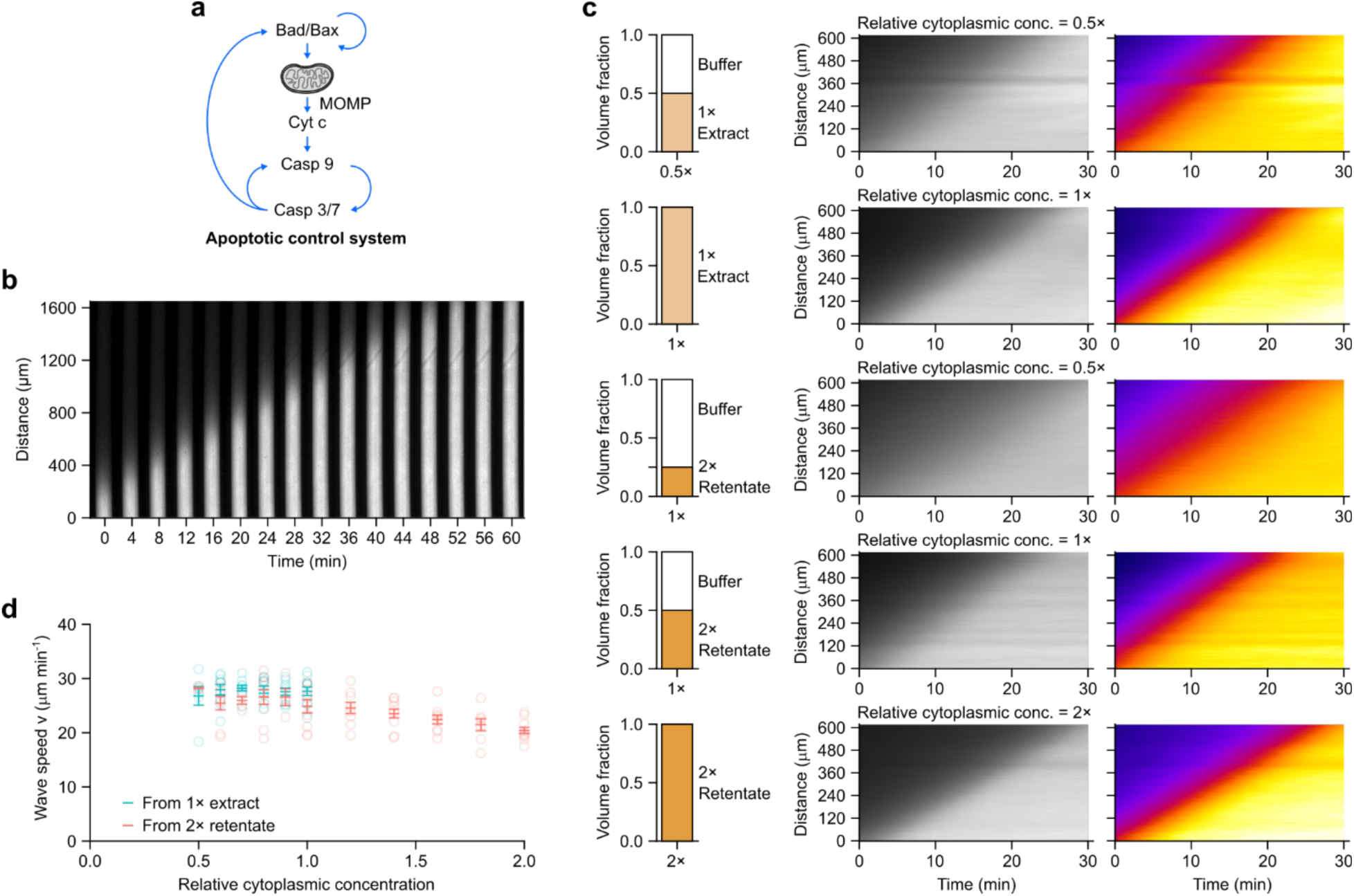
Apoptotic trigger wave speed is robust to change in cytoplasmic concentration. **a** Schematic view of the apoptotic control system. Note the multiple positive feedback loops. **b** A representative montage of an apoptotic trigger wave in a PTFE tube induced from the lower end. Bright signal is rhodamine 110 released by caspase 3/7 cleavage of (Z-DEVD)_2_-R110, which reports the activation of caspase 3/7. **c** Representative kymographs (middle and right columns) of extracts at different cytoplasmic concentrations prepared from either 1x extract (top 2 rows) or retentate (bottom 3 rows). Rhodamine 110 fluorescence is shown in grey scale (middle column) or pseudo-color (right column). Pseudo-coloring demonstrates that a range of fluorescence thresholds would give very similar estimates of trigger wave velocity. **d** Speeds of apoptotic trigger waves at different cytoplasmic concentrations. Means ± S.E.M. are shown. Data is compiled from 9 independent experiments.

Interphase *Xenopus* egg extracts were prepared and were mixed with a rhodamine-based fluorogenic sensor of caspase 3/7 activation, (Z-DEVD)_2_-R110, and a proteasome inhibitor (MG-132), which decreased the background level of R110 fluorescence and hence improved the signal-to-noise ratio of the experiment. The extracts were then loaded into thin (∼100 µm diameter) PTFE tubes (Fig. 2b). Apoptosis was induced by briefly dipping one end of the tube into a reservoir of apoptotic extract, prepared by adding cytochrome c (2 µM) to fresh extract and incubating at room temperature for 30 min. The induced tubes were then immersed in heavy mineral oil in custom-made imaging chambers and imaged at 2 min intervals at room temperature.

Fig. 2b shows the results of a typical experiment. Apoptosis, as detected by bright R110 fluorescence, first initiated at the dipped end of the tube and then spread toward the other end. The propagation speed in this experiment was 29.6 µm min^-1^; average speeds from 25 independent experiments were 27.5 ± 0.8 µm min^-1^ (mean ± S.E.M.), This is similar to the speeds seen in the cycling extracts that underwent apoptosis in Fig. 1E (28.1 and 27.7 µm min-1) and agree well with previous reports^8^.

We then altered the cytoplasmic concentration of the egg extract by diluting either a 1x extract or a 2x retentate. We verified apoptotic wave speed responds similarly to 3 different diluents (Supplementary Fig. 3) and chose XB buffer without sucrose as the primary diluent for further experiments. Fig. 2c shows kymographs of R110 fluorescence as a function of time for the original 1x extract and a 0.5x obtained by dilution, and for a 2x extract and a 1x extract reconstituted from the 2x extract by dilution; Fig. 2d shows data from 9 independent experiments, including additional extract concentrations. Overall, the apoptotic wave speed was almost invariant (Fig. 2d). There was no measurable change over a concentration range of 0.5x to 1x, and the speed decreased by only about 15% as the concentration increased to 2x (Fig. 2d). Thus, apoptotic wave speed is highly robust to variations in cytoplasmic concentration.

### Deriving an expression for wave speed as a function of cytoplasmic concentration

To try to understand these trends, and in particular to understand how wave speed can be so insensitive to cytoplasmic concentration, we derived an expression for wave speed as a function of cytoplasmic concentration based on a simple model of a reaction-diffusion trigger wave system. We began by assuming that the complicated reaction schemes shown in Fig. 1a (for mitosis) and Fig. 2a (for apoptosis) can be approximated by simple one-species, bimolecular autocatalytic processes, as shown in Fig. 3a and in Eq. 1:

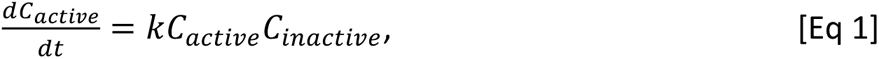

**Figure 3.**
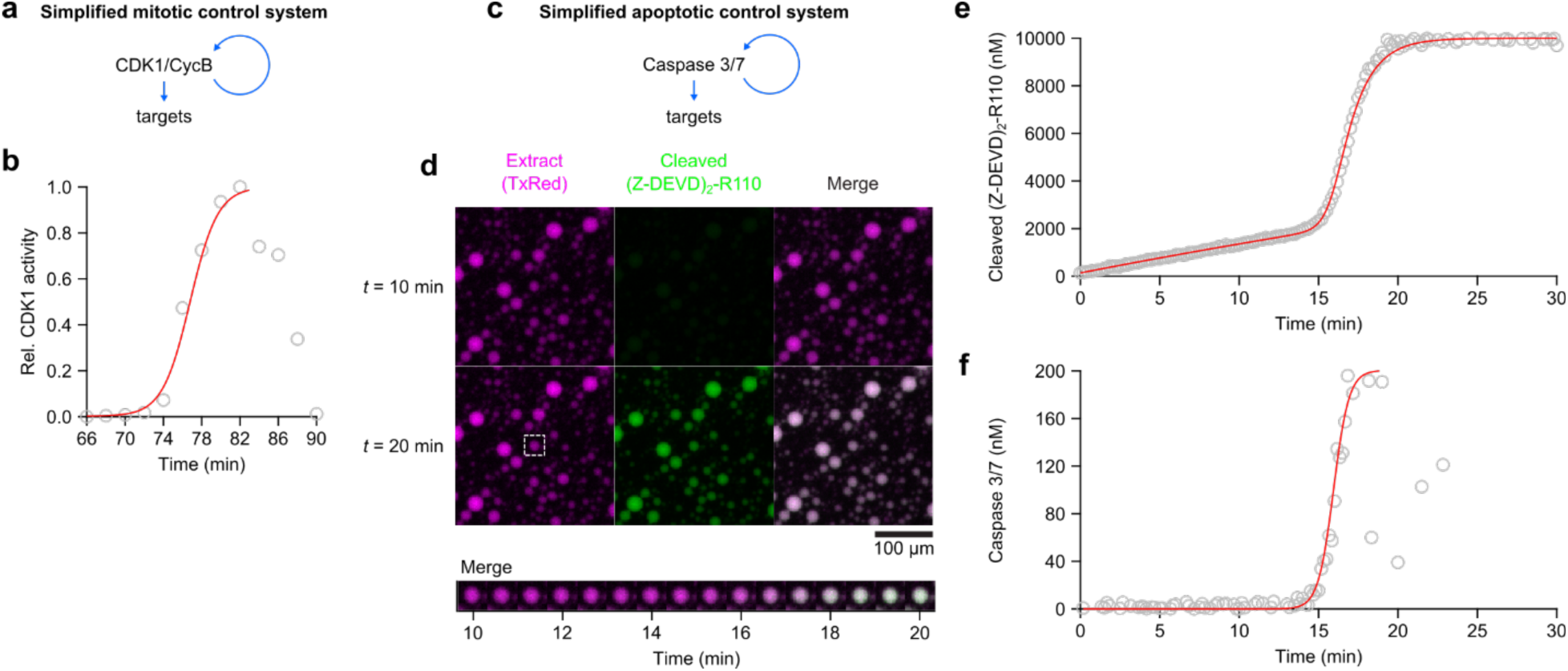
Mitotic and apoptotic activities are well-approximate by the logistic equation. **a** Mitotic control system can be conceptualized more simply as a single, combined positive feedback loop controlling CDK1/CycB at the onset of mitosis. **b** Relative CDK1 activity at mitotic onset, as measured by H1 kinase activity assay, can be well-fitted by a logistic model. Data are taken from Pomerening et al^36^. **c** The apoptotic control system can also be conceptualized as a single, combined positive feedback loop onto caspase 3/7. **d** Apoptosis in droplets of encapsulated extract. Extract containing added (Z-DEVD)_2_-R110 plus apoptotic extract and a cytoplasmic marker (TXRed) were encapsulate in squalene plus 5% (v/v) Cithrol DPHS-SO-(AP). Fluorescence was followed as a function of time by microscopy. **e** Kinetics of (Z-DEVD)_2_-R110 cleavage. Open circles are data at each time point. The red curve is based on the ODE model for caspase 3/7 activation and (Z-DEVD)_2_-R110 cleavage (Eqs 9 and 10) fitted to the data. **f** Kinetics of caspase 3/7 activation. Open circles are concentrations of active caspase 3/7 calculated based on the ODE model (Eqs 9 and 10). Red curve is the logistic growth curve fitted to the data.

where 𝐶*_𝑎𝑐𝑡*i*𝑣𝑒_* denotes the concentration of active Cdk1 or caspase 3/7, 𝐶*_*i*𝑛𝑎𝑐𝑡*i*𝑣𝑒_* is the concentration of inactive Cdk1 or caspase 3/7, and 𝑘 is a rate constant. We can eliminate one of the time-dependent variables by assuming that the total concentration of Cdk1 or caspase 3/7 is constant, and substituting as follows:

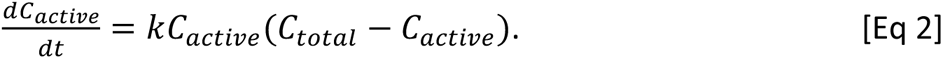

This ordinary differential equation can be solved in closed form:

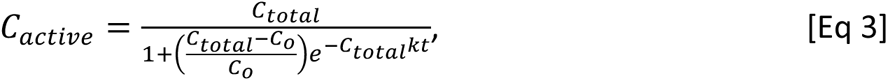

where 𝐶_0_ is 𝐶*_𝑎𝑐𝑡*i*𝑣𝑒_*(𝑡 = 0). Eq. 3 is a logistic equation, identical in form to the equation that describes the growth of bacteria in the face of limited resources. The time course is a sigmoidal curve, where the steepness of the curve is determined by 𝐶_𝑡𝑜𝑡𝑎𝑙_𝑘 and the time lag is determined by 𝐶_𝑡𝑜𝑡𝑎𝑙_, 𝐶_0_, and 𝑘.

To test whether it was reasonable to be replacing the complicated reaction schemes for Cdk1 and caspase 3/7 control with the simple rate equation shown in Eq. 2, we asked whether Eq. 3 could be fitted to time course data for Cdk1 activation and caspase 3/7 activation in extracts. For Cdk1 activation (Fig. 3a), we made use of previously published data on the time course of Cdk1 activity, measured by H1 kinase assays, in *Xenopus* extracts^36^. As shown in Fig. 3b, the time course was well approximated by a logistic function until Cdk1 activity began dropping during late mitosis. Cdk1 activity fell after reaching its maximum because of the activation of the APC/C and the destruction of cyclins, which causes the cycling system to exit mitosis.

To measure the time course of caspase 3/7 activation, we encapsulated a mixture of 10% apoptotic extracts and 90% fresh interphase extract, plus (Z-DEVD)_2_-R110, in squalene^37^, and monitored the increase in R110 fluorescence as a function of time in individual droplets (10 – 100 µm diameter; Fig. 3c, d). The time course for one such droplet is shown in Fig. 3e. The time course of caspase 3/7 activation could be inferred by assuming irreversible production of R110 fluorescence by active caspase 3/7 and a nonspecific background process. As shown in Fig. 3f, the time course was well-approximated by a logistic function. Estimated caspase activities fell after reaching the maximum due to depletion of the fluorogenic substrate.

The fact that the logistic function describes the early time course justifies the use of Eq. 2 to replace the complex, multivariable reactions of the full system. This makes our reaction-diffusion model equivalent to the Fisher-Kolmogorov-Petrovsky-Piskunov (FKPP) model^38,39^, which was originally formulated to describe the spreading of favorable genetic alleles through a population in space and time. For the one-dimensional case of trigger wave propagation in a thin tube, the resulting equation is:

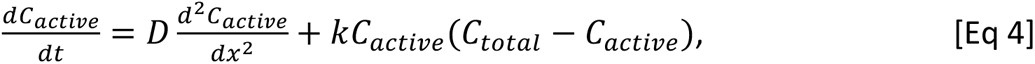

where 𝐷 is the diffusion coefficient for the autocatalytic species.

In the FKPP model, the minimum speed of propagation, which is the trigger wave speed, is given by:

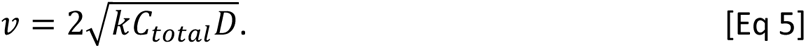

This is equivalent to Luther’s equation, proposed in 1906 to account for the speed of chemical waves^40,41^.

Note that all three of the variables under the square root sign might vary with the overall cytoplasmic concentration 𝜙. This can be expressed as:

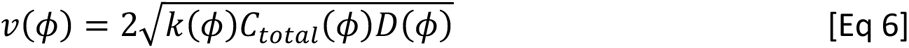

We assume 𝐶_𝑡𝑜𝑡𝑎𝑙_(𝜙) is simply proportional to the overall cytoplasmic concentration 𝜙:

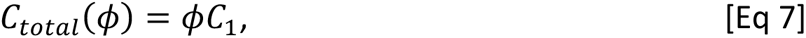

where 𝐶_1_denotes total caspase 3/7 or Cdk1 concentration at 1x cytoplasmic concentration. This assumption leaves two functions, (𝜙) and 𝑘(𝜙), to be experimentally determined.

For mitotic trigger waves, likely mediators of the spatial spread include cyclin B-Cdk1, Cdc25, and Gwl, proteins with molecular weights of ∼100 kDa. For apoptosis, plausible mediators include activated caspase 3 and 7 heterotetramers (∼60 kDa) and cytochrome c (12 kDa). Therefore, we chose a similarly-sized probe, Alexa Fluor 488-labeled bovine serum albumin (AF488-BSA; ∼67 kDa), for diffusion measurements. We measured its diffusion coefficient as a function of cytoplasmic concentration by fluorescence correlation spectroscopy (FCS). The diffusion mode of AF488-BSA was fairly close to Brownian in XB buffer without sucrose (α ≈ 0.9) and more subdiffusive in extracts (α ≤ 0.8; Fig. 4a, Supplementary Fig. 4). These observations are consistent with previous reports^42–45^. The effective diffusion coefficient 𝐷*_eff_*(𝜙), which is calculated assuming Brownian motion rather than subdiffusive motion, decreased exponentially with cytoplasmic concentration (Fig. 4b) (*R*^2^ = 0.953), as predicted by Phillies’ law^46,47^. The fitted effective diffusion coefficients in XB buffer without sucrose (𝜙 = 0) and 1x extract (𝜙 = 1) were 32 and 15 µm^2^ s^-1^ (Fig. 3b), respectively, again consistent with previous measurements^42^. We can therefore express the scaling of the effective diffusion coefficient 𝐷*_eff_*(𝜙) as:

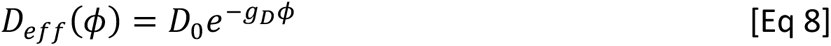

**Figure 4.**
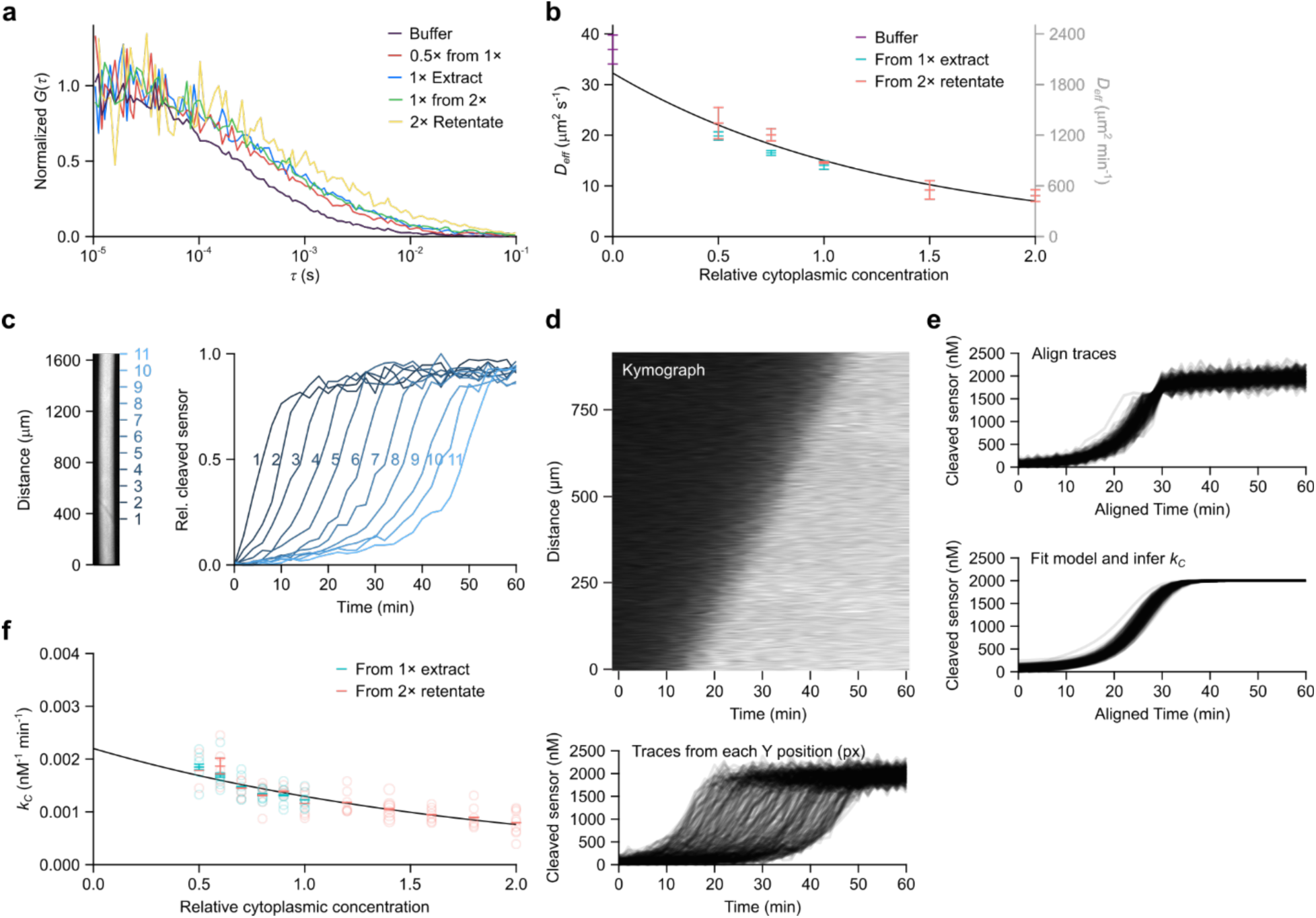
The effective diffusion coefficient of AF488-BSA and apparent autocatalytic rate constant of caspase 3/7 decrease exponentially with cytoplasmic concentration. **a** Representative fluorescence correlation spectroscopy (FCS) autocorrelation functions for AF488-BSA in extracts with different cytoplasmic concentrations. (𝑟) is the autocorrelation function and 𝑟 is the time delay. **b** Effective diffusion coefficient of AF488-BSA at different cytoplasmic concentrations. Effective diffusion coefficients were calculated by fitting the autocorrelation data from FCS measurements to a Brownian diffusion model. A 60 s fluorescence intensity time course was registered for each FCS measurement. Means ± 90% CI calculated from 3 measurements are shown. Solid black curve is an exponential curve fitted to the means. **c** (Z-DEVD)_2_-R110 cleavage kinetics can be monitored as apoptotic trigger waves sweep through a tube of extract. In this example, fluorescence from cleaved (Z-DEVD)_2_-R110 at 11 positions (left) are shown on the right. Fluorescence from the cleaved (Z-DEVD)_2_-R110 was normalized to the maximal value at each location. **d** (Z-DEVD)_2_-R110 cleavage kinetics in a kymograph (upper) can be represented as a series of traces (lower). **e** Traces shown in (**d**) were aligned by time (upper) and the model for caspase 3/7 activation and (Z-DEVD)_2_-R110 cleavage was fitted to the data. Individual fitted traces are shown in the bottom panel. **f** The apparent autocatalytic rate constant 𝑘_𝐶_ at different cytoplasmic concentrations was extracted from the fitted model. Shown are means ± S.E.M. compiled from the same 9 experiments as the ones shown in Fig. 2d. The black solid curve is an exponential curve fitted to the means.

where 𝐷_0_ is 𝐷*_𝑒ff_*(0) and 𝑔_𝐷_ is a dimensionless scaling factor with a fitted value of 0.765.

The remaining contributor to Eq 6 is (𝜙), the rate constant for the bimolecular autocatalysis reaction. We do not have a good way of experimentally assessing this parameter for Cdk1 activation, since we are inferring Cdk1 activity indirectly from microtubule polymerization, but the more direct probes for caspase activation allow this relationship to be determined. To infer enzyme activities from fluorescence data, we first constructed an ordinary differential equation (ODE) model for how (Z-DEVD)_2_-R110 dynamics depend upon caspase activity. We assumed that fluorescent rhodamine 110, designated 𝑅, is produced by caspase 3/7 and degraded by some unspecified enzyme, and that both production and degradation are approximated by mass action kinetics:

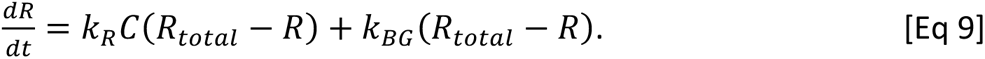

Here 𝑅_𝑡𝑜𝑡𝑎𝑙_ is the total concentration of (Z-DEVD)_2_-R110 and its fluorescent product R110, 𝑘_𝑅_ the rate constant for caspase 3/7 cleaving (Z-DEVD)_2_-R110, and 𝑘_𝐵𝐺_ is the first-order rate coefficient for background/nonspecific production of R110. Eq 9 can be solved to obtain an expression for *R*(𝑡):

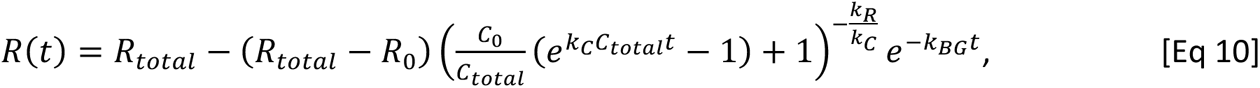

with 𝑅_0_ representing the fluorescent rhodamine 110 concentration and 𝐶_0_ the concentration of the apoptotic mediator at 𝑡 = 0. For clarity, we use 𝑘_𝐶_ to represent the bimolecular rate constant for caspase 3/7 autocatalysis, making it equivalent to 𝑘 in Eqs 4, 5, and 6.

There are 4 adjustable parameters in Eq 10; namely, 𝑘_𝐶_, 𝐶_0_, 𝑘_𝑅_, and 𝑘_𝐵𝐺_. We experimentally determined 𝑘_𝑅_ as a function of cytoplasmic concentration (Supplementary Fig. 5), thereby eliminating one adjustable parameter from Eq 10. We then experimentally measured the rate constant 𝑘_𝐶_ for the cleavage of (Z-DEVD)_2_-R110 by caspase 3/7 in *Xenopus* egg extracts by quantifying the sensor cleavage as a function of time at each point in the tube, aligning the time courses, and fitting the model to the aligned traces (Fig. 4d). This procedure was then repeated for various cytoplasmic concentrations.

**Figure 5.**
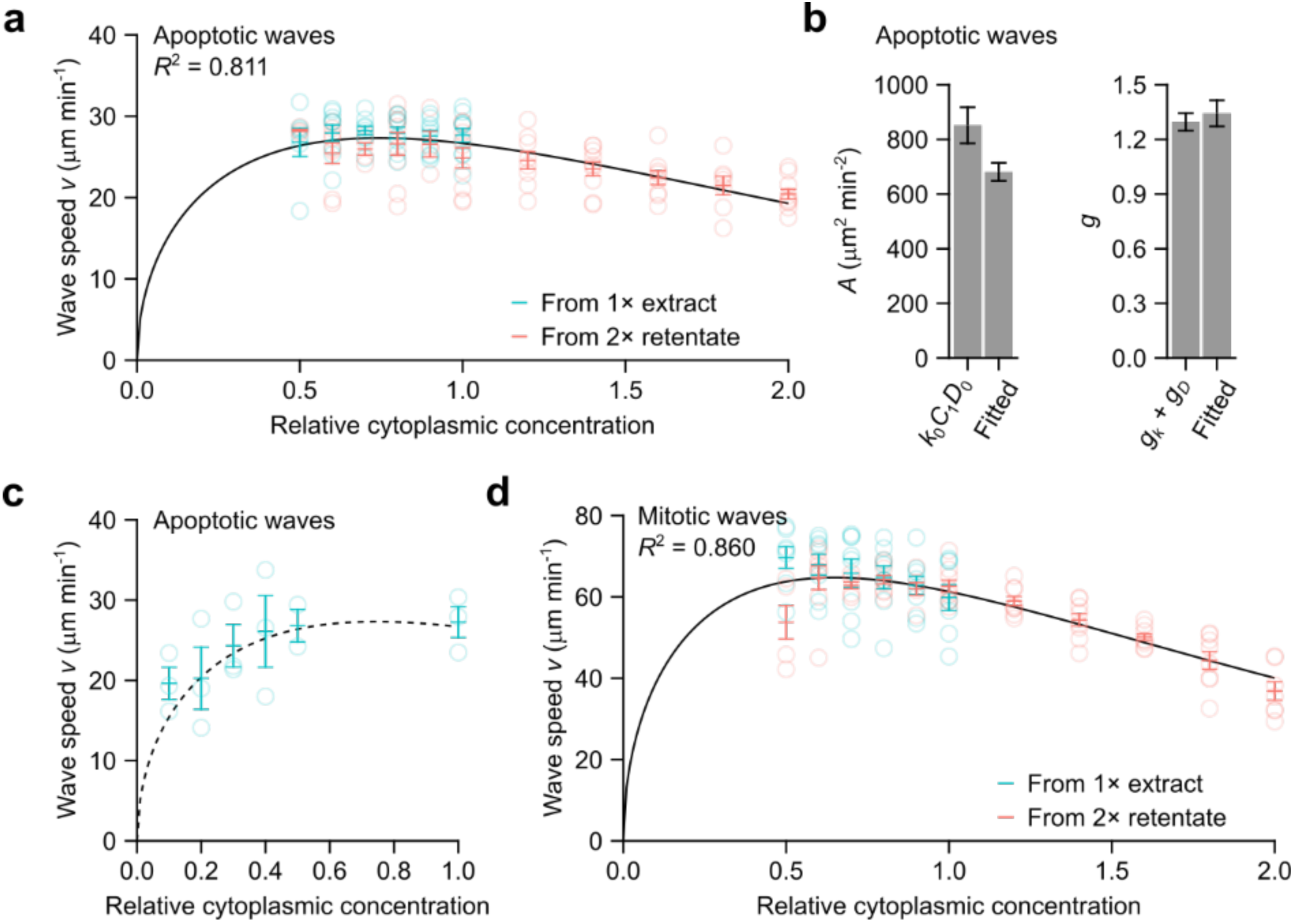
Mitotic and apoptotic trigger wave speeds at different cytoplasmic concentrations follow the generalized Luther’s equation. **a** Apoptotic wave speeds at different cytoplasmic concentrations as shown in Fig. 2D. The solid black curve is the generalized Luther’s equation fitted to the means. **b** The value of 𝐴, calculated as the product of the experimentally-determined parameters 𝑘_0_, 𝐷_0_, and 𝐶_1_, is compared to the fitted value (left panel). Likewise, the value of 𝑔 calculated as the sum of the experimentally-determined parameters 𝑔_𝑘_ and 𝑔_𝐷_, is compared to the fitted value (right panel). 𝑘_0_, 𝐷_0_, 𝑔_𝑘_, and 𝑔_𝐷_ are from the exponential fit shown in Fig. 4. Error bars are S.E.M.s calculated directly from the fittings or propagated from the individual experimentally-determined parameters. **c** Apoptotic wave speeds at low cytoplasmic concentrations. Dashed line shows the same fitted curve as in (**a**), which was obtained from only higher concentration data. Shown are means ± S.E.M. from 3 independent samples. **d** Mitotic wave speeds at different cytoplasmic concentrations, replotted from Fig. 1g and fitted to the generalized Luther’s equation (black curve).

One might expect 𝑘_*c*_(𝜙) to be roughly constant since most enzymes operate far from the calculated Smoluchowski limit for diffusion control. However, 𝑘_*c*_(𝜙) decreased exponentially with increasing cytoplasmic concentration (*R*^2^ = 0.901; Fig. 4e). We can therefore express the scaling of 𝑘_*c*_(𝜙) as:

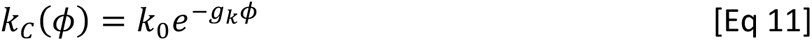

where 𝑘_0_ ≡ 𝑘_*c*_(𝜙 = 0) and 𝑔_𝑘_ is the scaling factor, which were empirically estimated to be of 0.0022 nM^-1^ min^-1^ and 0.531, respectively. We can then rewrite Luther’s equation to explicitly include the three concentration dependencies:

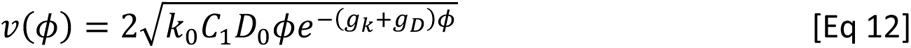

We can further simplify Eq 12 by defining:

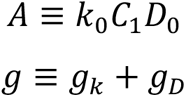

and arrive at the following expression:

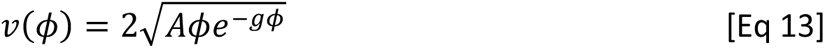

where 𝐴, the speed factor, determines the magnitude of the trigger wave speed, and 𝑔, the cytoplasmic concentration scaling factor, determines how steeply the speed decreases at high cytoplasmic concentration. Using *A* and *g* as adjustable parameters, Eq 13 fits well to the experimental data for apoptotic wave (Fig. 5a). We can also compare the fitted parameters to the quantities that contribute to them as estimated by experiments. Both 𝑔_𝑘_ and 𝑔_𝐷_were individually measured (Fig. 4b, f), and their sum is close to the fitted value of *g* (Fig. 5b). The product of the experimentally estimated values of 𝑘_0_, 𝐶_1_, and 𝐷_0_ was also in reasonable agreement with the fitted value (Fig. 5b). The agreement between the directly measured values for these parameters and the values inferred from the trigger wave speed measurements is reassuring.

Note that Eq 13 also predicts that at low cytoplasmic concentrations the wave speed should decrease, a trend that was not apparent in the initial experimental data (Fig. 5a). To test this prediction, we repeated the experiment over very low cytoplasmic concentrations, and, as shown in Fig. 5c, the low concentration data agreed well with curve fitting carried out on the higher concentration (0.5x to 2x) results alone. Taken together, these findings show that combining the FKPP expression for trigger wave speed and Phillies’ equation for the concentration dependence of diffusion-limited enzyme activities yields an equation that accounts for the dependence of trigger wave speed on cytoplasmic concentration, including the near-maximal speed at 1x concentration, the robustness of the trigger wave speed over a wide range of cytoplasmic concentrations, and the fall-off in speed at very high and very low concentrations.

Eq 13 could also be fitted well to the mitotic wave speed data (Fig. 5d). The fitted 𝑔 value was larger (1.54 vs. 1.34), which accounts for the observation that the wave speed fell more steeply with increasing cytoplasmic concentration.

### Mechanism of the robust apoptotic trigger wave speed

The robustness of the trigger wave speed appears to arise because one factor that influences wave speed, the concentration of the diffusible apoptotic mediator 𝐶_𝑡𝑜𝑡𝑎𝑙_, increases with increasing cytoplasmic concentration, whereas the autocatalytic rate constant 𝑘_𝐶_ and diffusivity 𝐷_𝑒*ff*_ decrease. Perfect robustness would arise if the competing trends canceled exactly. Here we examine in more detail how close to exact the cancelation is and why it breaks down at very high and very low cytoplasmic concentrations.

We first expressed these functions in relative terms:

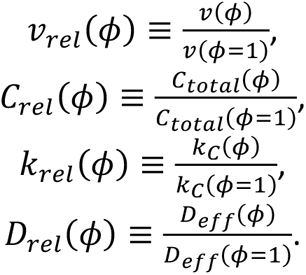

Defining these relative quantities allows us to focus on the dependencies on cytoplasmic concentration 𝜙 while the proportionality constants 𝐶_1_, 𝑘_0_, and 𝐷_0_ cancel out. We can also express Eq 6 in relative terms:

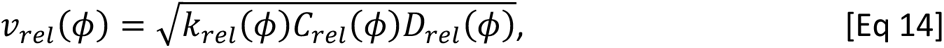

If we take the logarithm, then the individual factors combine additively rather than multiplicatively:

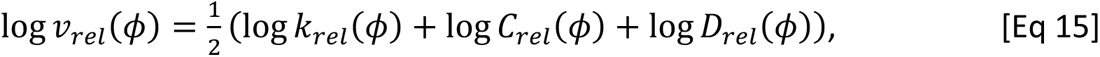

A robust trigger wave speed means 𝑣_𝑟𝑒*l*_(𝜙) should be close to 1 for a range of cytoplasmic concentration 𝜙. With log-transformation applied, a robust speed should have log 𝑣_𝑟𝑒*l*_(𝜙) close to 0, meaning that the right-hand side of Eq 15 should also be close to 0:

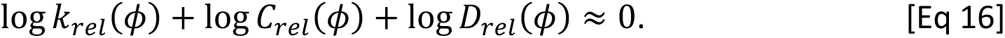

From 0.5x to 1x cytoplasmic concentration, the decrease in log 𝐷_𝑟𝑒𝑙_(𝜙) and log 𝑘_𝑟𝑒𝑙_(𝜙) combined (blue and green bars, respectively) is nearly equal to the gain in log 𝐶_𝑟𝑒𝑙_(𝜙) (red bars, plotted as negative to aid visual comparison; Fig. 6a). This explains why the apoptotic wave speed is almost constant over that range of concentrations. At higher cytoplasmic concentrations, the negative factors (log 𝐷_𝑟𝑒_(𝜙) and log 𝑘_𝑟𝑒𝑙_(𝜙)) are larger in magnitude than the positive factor (log 𝐶_𝑟𝑒𝑙_(𝜙)), and so the wave speed decreases with increasing cytoplasmic concentration, and at very low cytoplasmic concentrations, the opposite is true (Fig. 6a). Thus, the robustness of the trigger wave speed arises from the precise balancing of opposing kinetic and biophysical quantities over a range of cytoplasmic concentrations.

**Figure 6.**
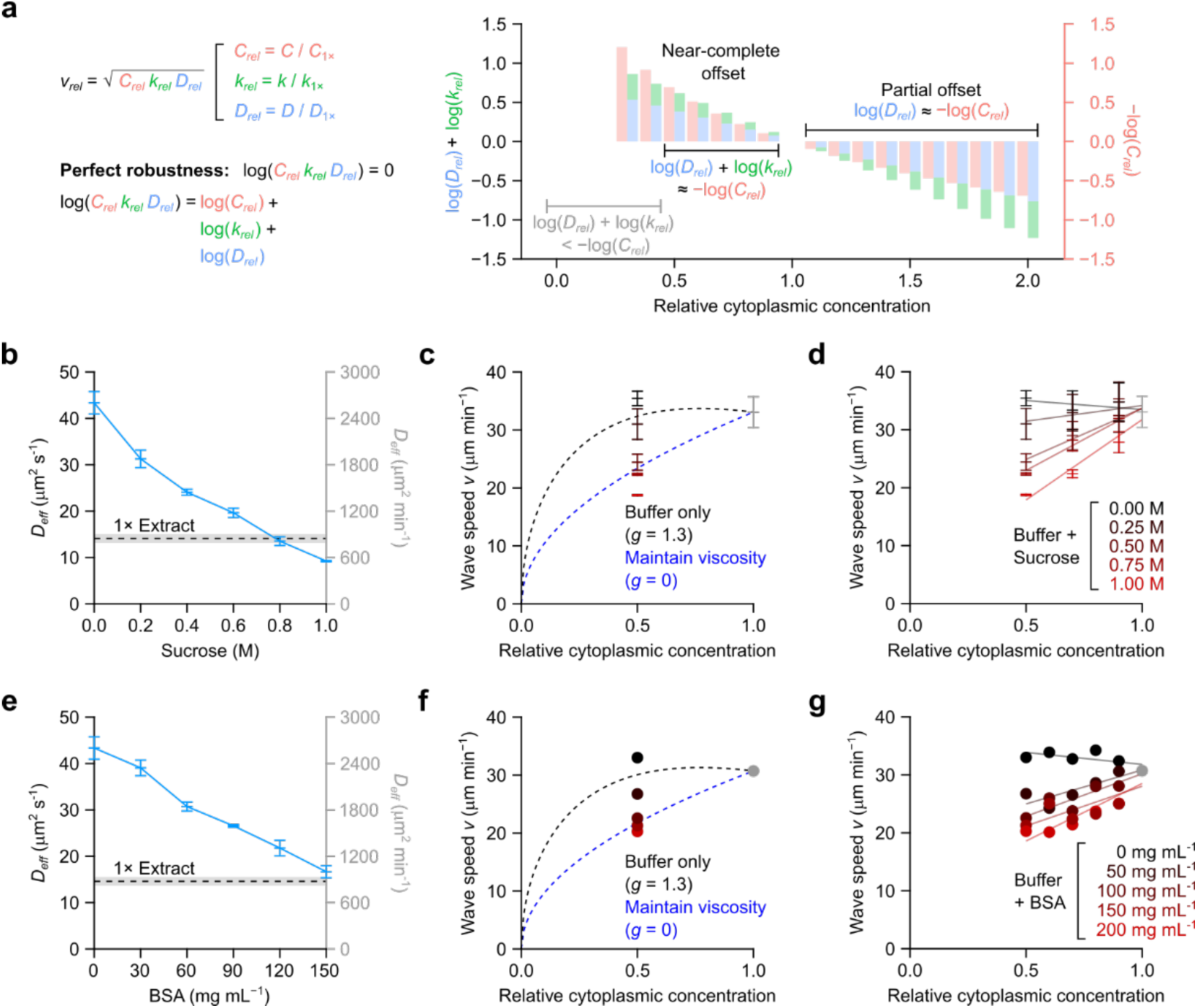
Opposing kinetic and physical effects give rise to robustness in trigger wave speeds. **a** Comparing the effect sizes of changes in rate constant, effective diffusion coefficient, and concentration. Effect size is defined as fold change relative to 1x cytoplasmic concentration. For simplicity, the data for cytoplasmic concentrations < 0.3x, where the concentration effect dominates, are omitted. **b**, **e** Effective diffusion coefficients of AF488-BSA in XB buffers of various sucrose concentrations (**b**; no BSA present) or BSA concentrations (**e**; no sucrose present) as determined by FCS. AF488-BSA diffusion in 1x extract can be mimicked by ∼0.8 M sucrose or ∼150 mg mL^-1^ BSA. **c**, **f** Apoptotic trigger wave speeds with extracts diluted with buffer (black data points) or a viscogen (red data points), and compared with theoretical curves (dashed lines). Crowding effects are quantified by the parameter 𝑔. For apoptotic trigger waves, 𝑔 is ∼1.3 (Fig. 5b) and is 0 if crowding effects are absent. We note that, in the case of 𝑔 = 0, wave speed 𝑣 follows the square root of total caspase 3/7 concentration. 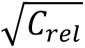. Means ± S.E.M. (n = 3) are shown for sucrose-containing buffers (**c**). Means (n = 2) are shown for BSA-containing buffers (**f**). The curves were set to pass through wave speeds at 1x cytoplasmic concentration for both sucrose-containing and BSA-containing buffers. Apoptotic wave speeds at 0.5x cytoplasmic concentration are also plotted. We note the scaling can be approximated by a horizontal line for 𝑔 = 1.3 between 0.5x and 1x cytoplasmic extract, whereas for 𝑔 = 0, a straight line with a positive slope. **d**, **g** Apoptotic trigger wave speeds are plotted for a range of sucrose-containing (**d**) or BSA-containing buffers (**g**) at different cytoplasmic concentrations. Means ± S.E.M. (n = 3) are shown for sucrose-containing buffers (**d**) whereas means (n = 2) are shown for BSA-containing buffers (**g**). Straight lines were fitted to each sucrose (**d**) or BSA (**g**) concentration. Only buffer without sucrose or BSA manifest straight lines with slightly negative slopes. Viscogen-containing buffers, be it sucrose or BSA, manifest positive slopes.

### Artificially maintaining diffusivity abrogates trigger wave speed robustness

One strong prediction is that if we could dilute an extract without increasing its diffusivity, the robustness of the trigger wave speed would be compromised. Toward this end we tested two viscogens, sucrose and BSA, and determined what concentrations would yield buffer solutions with diffusion coefficients equal to those seen in 1x cytoplasm. Using FCS and AF488-BSA, we found that 0.8 M sucrose (Fig. 6b) and 150 mg mL^-1^ BSA (Fig. 6e) yielded diffusivities equivalent to that of 1x cytoplasm. We then diluted 1x cytoplasmic extracts with these buffers and asked whether trigger wave speed was no longer robustly maintained. As shown in Fig. 6c, f, trigger wave speed was now dependent upon cytoplasmic concentration over this range. Intermediate concentrations of the viscogens yielded intermediate wave speed results (Fig. 6d, g). Thus the exact balancing of the effects of cytoplasmic diffusivity and reactant concentration is the basis for the robustness in trigger wave speed.

## DISCUSSION

### The robustness of trigger wave speeds

Over the past several years it has become increasing clear that trigger waves are a recurring theme in both intracellular^8,12^ and intercellular^13–18^ communication. Unlike diffusive spread, trigger waves allow signals to propagate without diminishing in amplitude or slowing in speed. Trigger waves are complex, systems-level phenomena; they require biological reactions that include positive feedback loops, plus a spatial coupling mechanism. Any system as complicated as this is bound to have vulnerabilities. Here we have examined how vulnerable two intracellular trigger waves, apoptotic waves and mitotic waves, are to variation in the cytoplasmic concentration, a basic cellular property that differs from cell type to cell type, and even varies in individual cells as they proceed through mitosis. We found that even though a priori one might expect that a bimolecular reaction’s speed would vary as the square of the cytoplasmic concentration, both apoptotic and mitotic wave speeds were nearly constant when extracts were diluted down from 1x to lower concentrations, and fell modestly at higher-than-physiological concentrations. We derived a simple model that accounts for both the robustness of trigger wave speed and the slowing seen at very high and very low cytoplasmic concentrations. The model implies that the robustness arises from canceling effects of cytoplasmic concentration: increasing cytoplasmic concentration should increase the speed by increasing the concentrations of the reactants, but should also decrease the speed by increasing viscosity and hence slowing both the local coupling process (diffusion) and the rate constants for the positive feedback reactions. This implies that if one were to change cytoplasmic concentration without changing viscosity, by diluting cytoplasm with buffer supplemented with the appropriate concentration of a viscogen, the wave speed should cease to be invariant, and indeed this was found to be the case. For large cells, such as the *Xenopus* eggs, the robustness of the mitotic wave speed could contribute to the reliability of the extremely rapid embryonic cell cycles in face of the physical stresses expected when an oocyte proceeds from the isotonic environment of the ovary to the hypotonic environment of the pond.

As mentioned above, recently it has been shown that some biological signals propagate as intercellular trigger waves in cell culture and in living tissues. Our theoretical framework may apply to tissue-level signal relay with appropriate generalization, with an intercellular process taking the place of intracellular diffusion as the local coupling mechanism.

This work adds to our burgeoning understanding of how physical properties of the cytoplasm constrain the operation of fundamental cellular processes. The present work also highlights the power of the *Xenopus* egg extract for studying the emergent regulatory functions that come from the differential responses of complex, coupled physical and biochemical processes.

### Limitations of the study

We relied on several approximations to access the analytical power of Luther’s and FKPP equations. Experimentally, we approximated diffusivity for caspase 3/7 and mitotic machines (CDK1/CycB and PP2A/B55 complexes) with AF488-BSA. Based on size, we may slightly over- or underestimate the scaling factor 𝑔_𝐷_, respectively. For diffusivity, we approximated anomalous diffusion with effective Brownian diffusion at a short distance range. Depending on the true length scale of diffusive mixing in trigger wave propagation, we may slightly over- or underestimate protein mobility. Model-wise, we approximated the overcoming of bistable switches (both mitotic and apoptotic onset are considered bistable transitions) as logistic growth processes. We may slightly underestimate the true rate coefficient depending on the true magnitude of ultrasensitivity. By adopting a combined positive feedback loop approach, we were not able to resolve the exact steps at which apoptosis or mitosis are limited by molecular crowding. Despite these limitations, our prediction error for apoptotic trigger wave speed was around 20% from the measured values, suggesting a good overall approximation.

Better experimental approximation and further developments in the theoretical treatments for anomalous diffusion and traveling waves in bistable media should improve the prediction accuracy and provide more detailed understandings to this topic.

## METHODS

### Key resource table

| **Chemicals, Peptides, and Recombinant Proteins** | **Source** | **Identifier** |
| --- | --- | --- |
| Pregnant mare serum gonadotropin | Fisher Scientific | 50-893-505 |
| Chorionic gonadotropin human | Sigma | CG10-10VL |
| L-cysteine | Sigma | 168149 |
| A23187 | Sigma | C7522 |
| Leupeptin | Sigma | L2023 |
| Chymostatin | Sigma | C7268 |
| Pepstatin A | Sigma | P5318 |
| Cytochalasin B | Sigma | C6762 |
| MG-132 | Selleckchem | S2619 |
| Cycloheximide | Sigma | 01810 |
| cOmplete protease inhibitor cocktail, EDTA-free | Sigma | 5892953001 |
| Sucrose | Sigma | S8501 |
| Bovine serum albumin | Sigma | A3294 |
| SiR-Tubulin | Cytoskeleton, Inc. | CY-SC002 |
| (Z-DEVD)<sub>2</sub>-R110 | AAT Bioquest | 13430 |
| Alexa Fluor 488 labeled bovine serum albumin | Thermo Fisher | A13100 |
| Cytochrome c purified from equine heart | Sigma | C2867 |
| **Experimental model: Organisms/strains** | **Source** | **Identifier** |
| *Xenopus laevis* females and eggs | NASCO | LM00531 |
| *Xenopus laevis* males and sperms | NASCO | LM00715 |
| **Reagents and materials** |  |  |
| Mineral oil, heavy | Sigma | 330760 |
| PTFE tubes | Cole Parmer | EW-06417-72 |
| 10 kDa molecular weight cut-off filter | Sigma | UFC501096 |
| Bio-Rad protein assay dye concentrate | Bio-Rad | 5000006 |
| SimplyBlue SafeStain | Thermo Fisher | LC6065 |
| **Software and Algorithms** | **Source** | **Identifier** |
| ZEN | Zeiss | N/A |
| Igor Pro, version 6 | WaveMetrics | https://www.wavemetrics.com/ |
| FIJI / ImageJ, | FIJI | https://imagej.net/software/fiji/ |
| R, version 4.0.2 | The R project | https://www.r-project.org/ |
| SAEMIX | CRAN | https://cran.r-project.org/web/packages/saemix/index.html |
| Mathematica, version 13.2 | Wolfram Research | https://www.wolfram.com/mathematica/ |

### Resource availability

#### Lead contact

Further information and requests for resources and reagents should be directed to and will be fulfilled by the lead contact.

#### Material availability

Materials used in this study will be made available upon request.

#### Data and code availability

Datasets and custom-written codes generated in this study will be made available upon request.

### Experimental model and subject details

#### Xenopus laevis

The animal work adhered to relevant national and international guidelines and received approval from the Stanford University Administrative Panel on Laboratory Animal Care (APLAC protocol 13307). *Xenopus laevis* females, aged over 3 years, were prepared by dorsal cavity injection of 100 U of pregnant mare serum gonadotropin (PMSG) at least 3 days prior to, but typically no more than 2 weeks before ovulation. Ovulation was induced through dorsal cavity injection of 200 U of human chorionic gonadotropin (hCG). These induced females were then housed in separate chambers containing egg-laying buffer (100 mM NaCl, 2 mM KCl, 1 mM MgSO_4_, 2.5 mM CaCl_2_, 500 µM HEPES, 100 µM EDTA, pH 7.4).

#### Preparation of demembranted *Xenopus* sperm chromatin

Demembranated *Xenopus laevis* sperm chromatin was prepared as described by Murray^48^. Typical experiments with cycling extracts included sperm nuclei, which serve as pacemakers for mitotic waves, at ∼100 nuclei per µL extract.

#### Preparation of *Xenopus* egg extracts

We followed previously established procedures^49^ to prepare *Xenopus* egg extracts with slight adaptations. Specifically, eggs were collected approximately 22 hours post hCG injection, selecting high-quality eggs with consistent pigment distribution and a well-defined white spot on the animal pole. The jelly coats were removed by incubation with dejellying buffer (20 mg mL^-1^ L-cysteine, pH 7.8) for no more than 5 min. Subsequently, the eggs underwent a minimum of three washes in 0.2× Marc’s modified Ringer’s solution (0.2× MMR; 20 mM NaCl, 400 µM KCl, 400 µM CaCl_2_, 200 µM MgCl_2_, 1 mM HEPES, 20 µM EDTA, pH 7.8).

For cycling extracts, the eggs were activated with calcium ionophore A23187 (0.5 µg mL^-1^) in 0.2× MMR prior to packing and crushing. The A23187-containing buffer was promptly removed upon egg activation, generally within 2 min, determined by the contraction of the animal pole. In the case of interphase-arrested extract, the activation step was omitted. After activation, the eggs were thoroughly washed with crushing buffer (50 mM sucrose, 100 mM KCl, 100 µM CaCl_2_, 1 mM MgCl_2_, 10 mM HEPES-KOH, pH 7.7) at least twice before packing through low-speed centrifugation (200 *g* for 1 minute, followed by 600 *g* for 30 s). After packing, care was taken to remove excess crushing buffer above the eggs to minimize dilution. Importantly, for cycling extracts, we waited at least 20 min post-activation to ensure meiotic exit was completed before transferring the packed eggs to ice, followed by subsequent centrifugation.

The eggs were subsequently crushed using a centrifugal force of 16,000 *g* at 4°C for 15 min. The resulting cytoplasmic fraction was collected as *Xenopus* egg extract and kept on ice. Peptidase inhibitor mix (10 µg mL^-1^ leupeptin, 10 µg mL^-1^ pepstatin, 10 µg mL^-1^ chymostatin) and actin polymerization inhibitor cytochalasin B (10 µg mL^-1^) were added into the extract. In the case of interphase-arrested extract, cycloheximide (CHX; 100 µg mL^-1^) was included to prevent cyclin B translation and, thus, entry into mitosis. CHX was excluded from cycling extracts. The extracts underwent one or two additional rounds of centrifugation at 16,000 *g* at 4°C for 5 min to eliminate impurities before further use.

#### Dilution, concentration, and reconstitution of *Xenopus* egg extracts

We used XB buffer without sucrose (100 mM KCl, 100 µM CaCl_2_, 1 mM MgCl_2_, 10 mM HEPES-KOH, pH 7.7) as the primary diluent for adjusting cytoplasmic concentration. In experiments involving viscogens, sucrose or BSA were titrated in XB buffer without sucrose to the required concentrations from stocks with the same salt content as the basal XB buffer. We concentrated the extracts using a 10 kDa molecular weight cut-off centrifugal filter. We could achieve a 2-fold concentration by centrifuging three times for 10 min at 4°C (a total of 30 min), homogenizing the extracts between each spin with gentle pipetting. Dilution and reconstitution were carried out by mixing extracts with the appropriate diluents to reach the desired volume fraction.

#### Determination of protein concentration

Protein concentration was determined using the Bio-Rad protein assay, a method based on the Bradford assay. Briefly, undiluted extract was first diluted 200-fold in XB buffer without sucrose and then quantified in accordance with the manufacturer’s instructions, measuring absorbance at 595 nm. The retentate was first diluted two-fold and quantified in the same manner as the undiluted extract. We also determined the protein concentration in the filtrate (flow-through) during the concentration process. The filtrate was directly assessed using the Bio-Rad assay, without additional dilution or processing.

### Measurements of apoptotic and mitotic trigger wave speeds

#### Experimental setup

To monitor apoptotic and mitotic trigger waves, we mixed extracts with biosensors and filled PTFE tubes (∼100 µm inner diameter) for subsequent time-lapse fluorescent microscopy. We used interphase-arrested extracts for apoptotic trigger wave propagation. Caspase 3/7 activity was monitored using (Z-DEVD)_2_-R110 at 2 µM, unless otherwise specified. To enhance the signal-to-noise ratio, we included 200 µM MG-132 to inhibit the proteasomal cleavage of (Z-DEVD)_2_-R110. Following tube filling, we let the extract-filled tubes stand at room temperature for 30 min before inducing apoptosis. Apoptosis was initiated by briefly dipping the tube end into a reservoir of apoptotic extract, which was prepared by adding 2 µM cytochrome c to fresh extract and incubating at room temperature for 30 min. The induced tubes were placed in custom-made imaging chambers, submerged in heavy mineral oil, and imaged at 2-min intervals at room temperature.

To monitor mitotic trigger waves, cycling extracts were used. We tracked the dissolution of microtubules during mitosis as an indicator of mitotic activity, with SiR-tubulin (200 nM) serving to visualize polymerized microtubules. Following the filling of PTFE tubes (∼100 or ∼300 µm in diameter), the tubes were immersed in heavy mineral oil within custom-made imaging chambers and imaged at 2 to 3-minute intervals at room temperature.

For each cytoplasmic concentration, we typically had several biological replicates (independent samples), which involved extracts prepared from different clutches of eggs obtained from different females. Within each biological replicate, several tubes (up to 5) were monitored as technical replicates. Typically, 1 to 2 waves per technical replicate were observed for apoptotic wave experiments, while more than 2 waves per technical replicate were typical for mitotic wave experiments.

#### Measurement of wave speeds from kymographs

Kymographs were constructed from time-lapse videos capturing the propagation of trigger waves within PTFE tubes using Multi Kymograph in FIJI/ImageJ, with the width parameter set to 3 pixels. The dimensions and signal intensity of the kymographs were adjusted to optimize visual inspection. To facilitate speed measurements of mitotic waves, we increased the contrast of the SiR-tubulin signal for better visualization. For apoptotic waves, images were binarized with a consistent signal cut-off by applying a unique global signal threshold to each image. Subsequently, straight lines were manually fitted to the linear segments of the propagating waves, and wave speed was determined from the slopes of these fitted lines.

Each biological replicate (independent sample) included up to 5 tubes as technical replicates. A median speed was initially calculated for each tube (tube median), followed by the determination of a median speed for a given biological replicate (biological replicate median) from the tube medians. Means and S.E.M. for each condition were then computed from the biological replicate medians.

### Measurement of protein diffusivity by FCS in *Xenopus* egg extract

FCS measurements in *Xenopus* egg extract were analyzed following a previously described method^42^. In brief, interphase-arrested extracts were prepared, and cytoplasmic concentration was adjusted as mentioned earlier. We included the EDTA-free cOmplete protease inhibitor cocktail at a 1:50 (v/v) ratio, 30 min prior to the addition of 25 nM Alexa Fluor 488 labeled BSA (AF488-BSA) to the extracts. FCS data were acquired using an inverted Zeiss LSM 780 multiphoton laser scanning confocal microscope at room temperature (22°C). The microscope setup and the calibration step were described previously^42^.

The confocal spot was focused 30 – 40 µm above the dish surface. Each data point represented the average of at least 3 randomly selected positions within the extract field. At each position, fluorescence intensities were acquired for 60 s. Autocorrelation functions were calculated directly by the ZEN 2.3 SP1 FP3 310 (Black) software (version 14) (Zeiss) controlling the microscope. An anomalous diffusion model or a Brownian diffusion model were used to fit the autocorrelation functions:

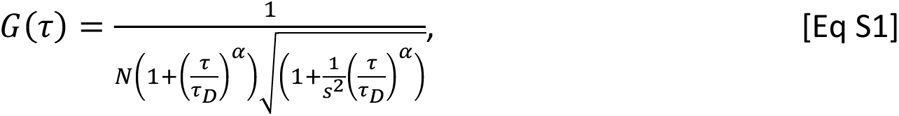

where 𝐺(*τ*) denotes the autocorrelation function, 𝛼 signifies the diffusion-mode parameter as defined by the mean squared displacement (MSD) equation 𝑀𝑆𝐷(𝑡) 𝖺 𝑡^𝛼^, *τ*_𝐷_ represents the characteristic diffusion time, 𝑁 corresponds to the particle number, and 𝑠 stands for the structural parameter of the optics. The Brownian diffusion model is identical to the anomalous model except for the 𝛼 value, which is set to 1.

Consistent with prior reports^42^, protein diffusion in *Xenopus* egg extract displayed weakly subdiffusive behavior, akin to cultured mammalian cells, and was better described by the anomalous diffusion model. Nevertheless, an effective diffusion coefficient can be calculated from 𝐷*_𝑒*ff*_* = 〈*τ*^2^〉/(4*r*_𝐷_) where *r* is the radius of the confocal spotsize.

### Logistic dynamics approximation of mitotic and apoptotic activities

#### CDK1 activity in cycling *Xenopus* egg extract

Measurements of CDK1 activity in the cycling *Xenopus* egg extract were made previously by Pomerening et al^36^. We focused on data points corresponding to the onset and exit of mitosis (60 to 90 min). The relative CDK1 activity was renormalized, and a logistic curve was fitted to the data within the selected time range up to the point when maximal activity was reached.

#### Caspase 3/7 activation in encapsulated extract droplets

For encapsulating interphase extract in oil, we followed the method outlined by Good and Heald^37^. In brief, we mixed 10% (v/v) apoptotic extract with fresh extract to induce the onset of apoptosis. Additionally, we included 10 µM (Z-DEVD)_2_-R110 to monitor caspase 3/7 activity. A low concentration of TexasRed-labeled dextran was added as a soluble marker for extract volume during image analysis. Subsequently, we added 1:20 volume of extract into squalene supplemented with 5% (v/v) Cithrol DPHS-SO-(AP) and agitated the tube with force to create an emulsion of encapsulated extract droplets. These droplets were then imaged at 10-second intervals using an epifluorescence microscope. The increase in fluorescence in the droplets was individually tracked using ImageJ.

We estimated active caspase 3/7 concentration over time by employing a set of ordinary differential equations (ODEs) as described in the main text (Eqs 2, 3, 9, and 10; see below). Essentially, caspase 3/7 activation was analyzed as an irreversible, bimolecular reaction following mass action principles. The cleavage of (Z-DEVD)_2_-R110 by caspase 3/7 was similarly modeled as an irreversible, bimolecular reaction based on mass action principles, while its nonspecific background cleavage in the extract was represented as an irreversible, first-order reaction following mass action.

### Estimation of caspase 3/7 activation rate coefficient

#### Model for caspase 3/7 activation and (Z-DEVD)_2_-R110 cleavage

We deduced caspase 3/7 activation kinetics from the cleavage kinetics of (Z-DEVD)2-R110. To do this, we constructed a simple model with two ordinary differential equations (ODEs) based on two key approximations. First, we approximated caspase 3/7 activation kinetics with logistic growth. Second, we approximated caspase 3/7 activation and (Z-DEVD)2-R110 cleavage at the pixel resolution using the following ODE:

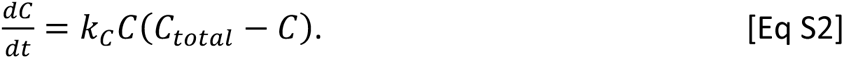

Here, 𝐶 denotes concentration of active caspase 3/7, 𝐶_𝑇𝑜𝑡𝑎𝑙_ the total concentration of inactive and active caspase 3/7, and 𝑘_𝐶_ the rate coefficient for the activation of caspase 3/7 through positive feedback. Logistic growth can be described in the following closed-form expression:

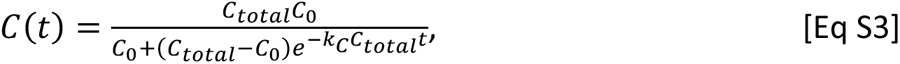

where 𝐶_0_ denotes caspase 3/7 concentration at 𝑡 = 0. Activated caspase 3/7 and nonspecific background activity consume (Z-DEVD)_2_-R110 and release fluorescent rhodamine 110 (R110):

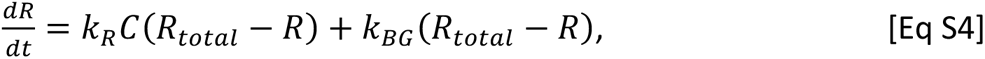

Here 𝑅 denotes free, fluorescent R110, 𝑅_𝑡𝑜𝑡𝑎𝑙_ the total concentration of (Z-DEVD)_2_-R110 and R110, 𝑘_𝑅_the rate coefficient of caspase 3/7 cleaving (Z-DEVD)_2_-R110, and 𝑘_𝐵𝐺_ the first-order rate coefficient of background cleavage. To reduce the number of fitting parameters in the equation, we experimentally determined 𝑘_𝑅_ for cytoplasmic concentration ranging from 0.5x to 2x (see below). We found a solution to (𝑡):

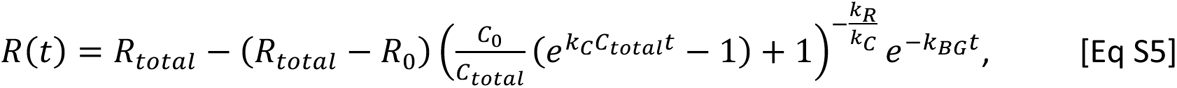

with 𝑅_0_ representing fluorescent R110 concentration at 𝑡 = 0.

#### Determination of the apparent (Z-DEVD)_2_-R110 cleavage rate coefficient

We measured (Z-DEVD)_2_-R110 cleavage rate in freshly prepared, fully apoptotic extracts. Briefly, apoptosis was induced by introducing 2 µM cytochrome c to interphase extracts and incubated for 30 min at room temperature to fully activate caspase 3/7. Subsequently, 5 µM (Z-DEVD)_2_-R110 was introduced to the apoptotic extracts, mixed vigorously by high-speed vortex, and the fluorescence increase was promptly recorded. Maximal cleavage rates were determined from the initial time points using linear fitting (*R*^2^ ≥ 0.99). The apparent second-order rate coefficients were then calculated from these maximal rates.

We made two assumptions. First, we assumed that caspase 3/7 concentration is linearly proportional to the overall cytoplasmic concentration and is 200 nM in 1x extracts. Second, we assumed a negligible decrease in (Z-DEVD)_2_-R110 during the early time points.

#### Estimation of (Z-DEVD)_2_-R110 cleavage kinetics from image brightness

The same kymographs employed for trigger wave speed measurements were utilized here. Signals were scaled, and the signal intensity was subsequently correlated with the nominal concentration of R110. To estimate the signal intensity asymptote, we employed a two-component Gaussian mixture model on the kymograph, with the larger component conveniently providing an estimate of the asymptote. Subsequently, we subtracted the background intensity and adjusted the signal accordingly. The signal intensity at 𝑡 = 0, therefore, also provided an estimate of 𝑅_0_, the initial rhodamine 110 concentration, reducing the parameters to be fitted to just three: 𝐶_0_, 𝑘_𝐶_, and 𝑘_𝐵𝐺_.

#### Model fitting

The fitting process was complicated by the spatial and temporal order of caspase 3/7 activation due to the propagation of apoptotic waves. To mitigate this effect, we aligned the trajectories of (Z-DEVD)_2_-R110 (with respect to time) and defined a relative time (also in units of minutes). We again utilized the two-component Gaussian mixture model fit and established a threshold at which the rhodamine 110 signal was 1000 times more likely to fall within the greater component than the lesser one. All trajectories were aligned to the first time point to pass this threshold, and a time window of 60 min around this time point was selected. The model described above was fitted to the R110 concentration within this time window. For this fitting, we employed the SAEMIX package in R, which is a stochastic approximation-based mixed-effect model approach. The SAEMIX algorithm is more robust and time-efficient than the commonly used nonlinear least square (NLS) algorithm for this purpose.

The mixed-effect model decomposes a “population” of varied observations into a population-level mean effect and individual-level random effects. We reported the population-level 𝑘_𝐶_ and only considered individual-level 𝑘_𝐶_ when demonstrating the individual-level fit. As in wave speed measurements, we calculated medians from observations within the same tube and then from tubes of the same biological replicate. The means and S.E.M.s were reported from all biological replicates.

#### List of parameters and variables

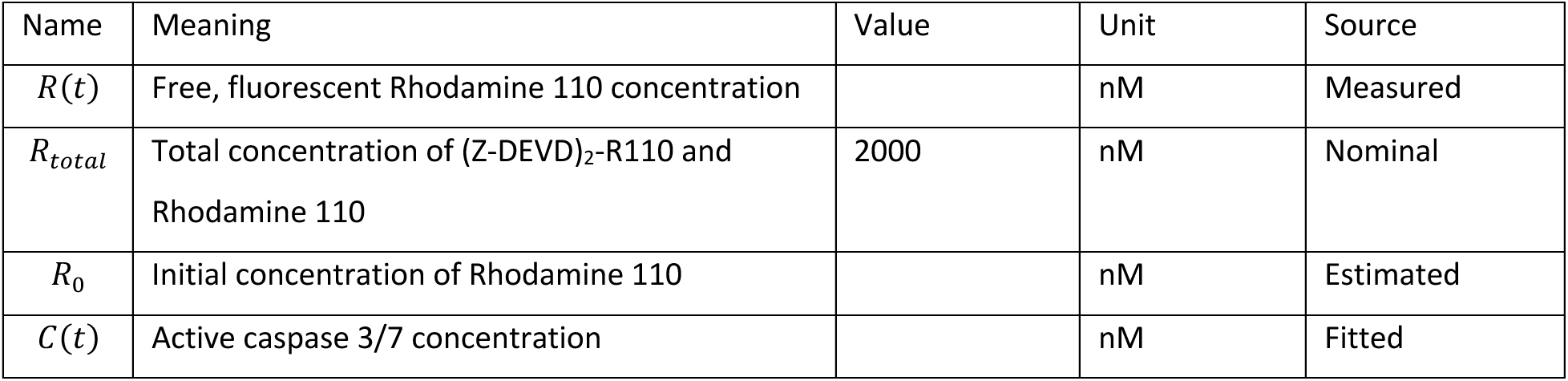
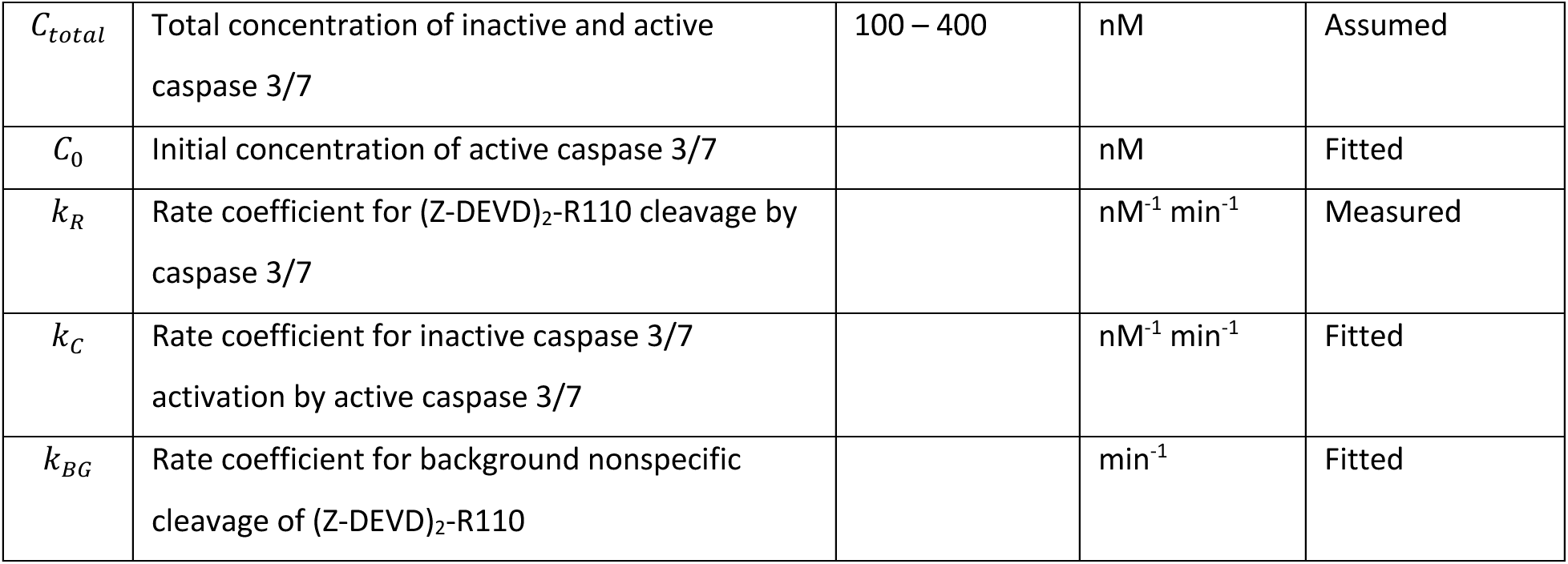

### Model fitting for scaling relationships

We fitted a linear model to the log-transformed means of the effective diffusion coefficient, 𝐷*_eff_*, and the rate coefficient, 𝑘_𝐶_. Additionally, we employed the generalized Luther’s equation to fit the means of trigger wave speed. In the case of the generalized Luther’s equation, fitting was carried out using the NLS algorithm within the stats package in R.

## Acknowledgements

This work was supported by a grant from the NIH (R35 GM131792) to J.E.F. and (K99 GM143481) to W.Y.C.H.

## Author contributions

J.H., Y.C., and J.E.F. conceptualized the study. J.H., Y.C., and S.T. performed the extract experiments. W.Y.C.H. and J.H. performed FCS experiments. J.H., Y. C., and J.E.F. analyzed the data. J.H. and J.E.F. conceptualized the model. J.H. and J.E.F. wrote the manuscript. J.E.F. supervised the study. J.E.F. and W.Y.C.H. secured the funding.

## Declaration of competing interests

The authors declare no competing interests.

---

**Supplementary Figure 1, related to Fig. 1.**
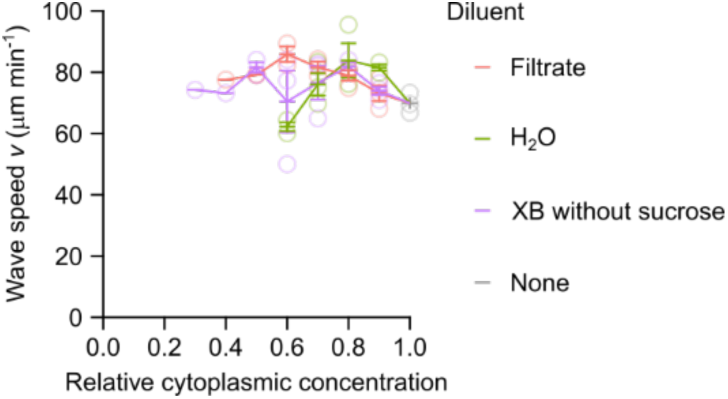
Mitotic trigger wave speed is robust to dilution using either filtrate or buffer. Speeds of mitotic trigger waves at different cytoplasmic concentrations. The cycling extracts were diluted using either filtrate, XB buffer without sucrose, or water. Note that dilution with filtrate and buffer produced comparable wave speeds, whereas dilution with water resulted in a drop in speed. For extracts diluted with water below 0.6x, cycle progression was not observed. Means ± S.E.M. compiled from 3 independent samples are shown.

**Supplementary Figure 2, related to Fig. 1.**
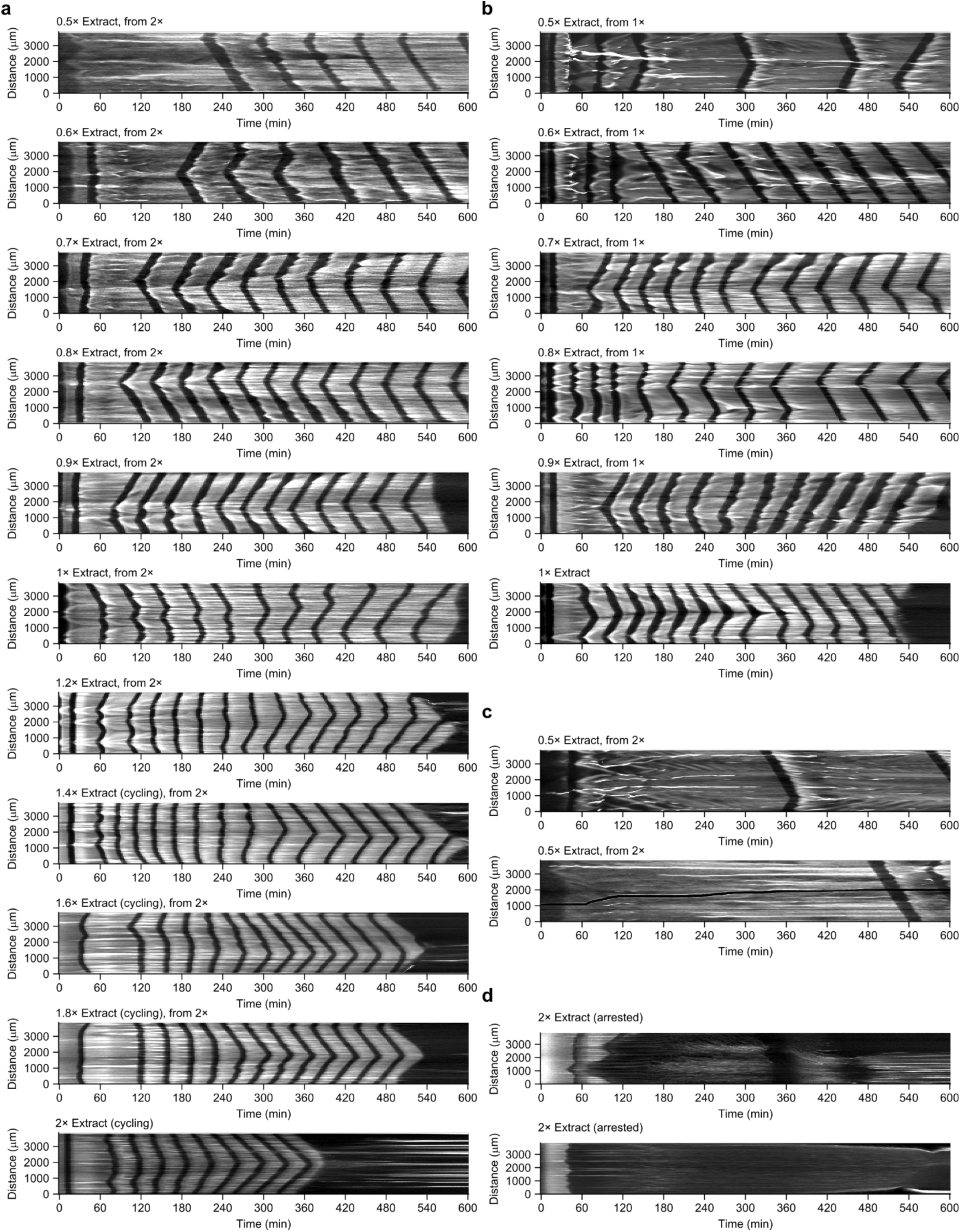
Cell cycle and mitotic trigger waves at different cytoplasmic concentrations. **a**, **b** Representative kymographs depict cell cycle and mitotic trigger waves at various cytoplasmic concentrations, prepared from 2x retentate (**a**) and 1x extract (**b**). **c** Two additional instances showcase 0.5x extracts prepared from 2x retentate, highlighting the variability in the first complete cell cycle across different extracts. **d** In contrast to the continuous cycling observed in Fig. 1e, these two examples of 2x retentate underwent mitotic arrest, occurring either in the first mitosis or the second.

**Supplementary Figure 3, related to Fig. 2.**
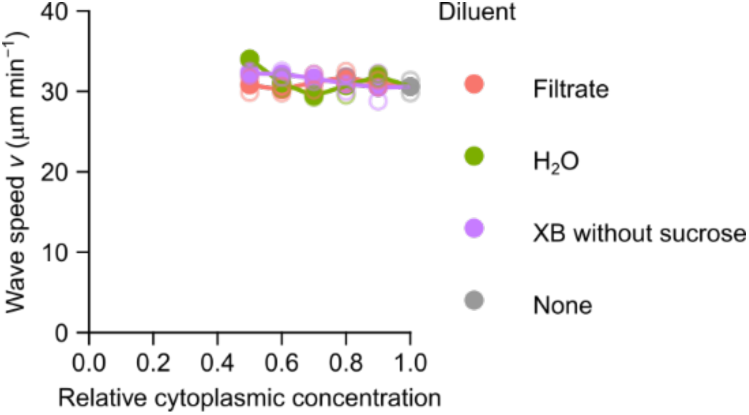
Apoptotic trigger wave speed is robust to dilution using filtrate, buffer, and water. Speeds of apoptotic trigger wave at different cytoplasmic concentrations. The interphase extracts were diluted using either filtrate, XB buffer without sucrose, or water. Means from 2 independent samples are shown.

**Supplementary Figure 4, related to Fig. 4.**
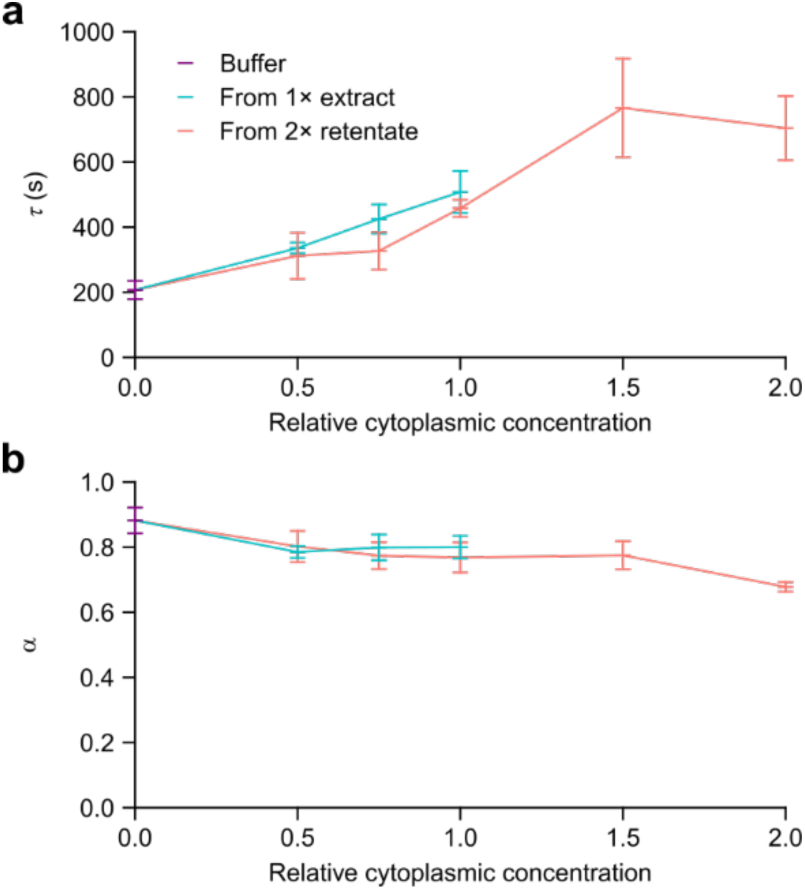
Anomalous diffusion fit to AF488-BSA diffusion in extracts. **a**, **b** FCS autocorrelation functions of AF488-BSA in extracts analyzed using the anomalous diffusion framework. Diffusion time *τ*_𝐷_ (**a**) and anomalous diffusion exponent 𝛼 (**b**) at different cytoplasmic concentrations.

**Supplementary Figure 5, related to Fig. 4.**
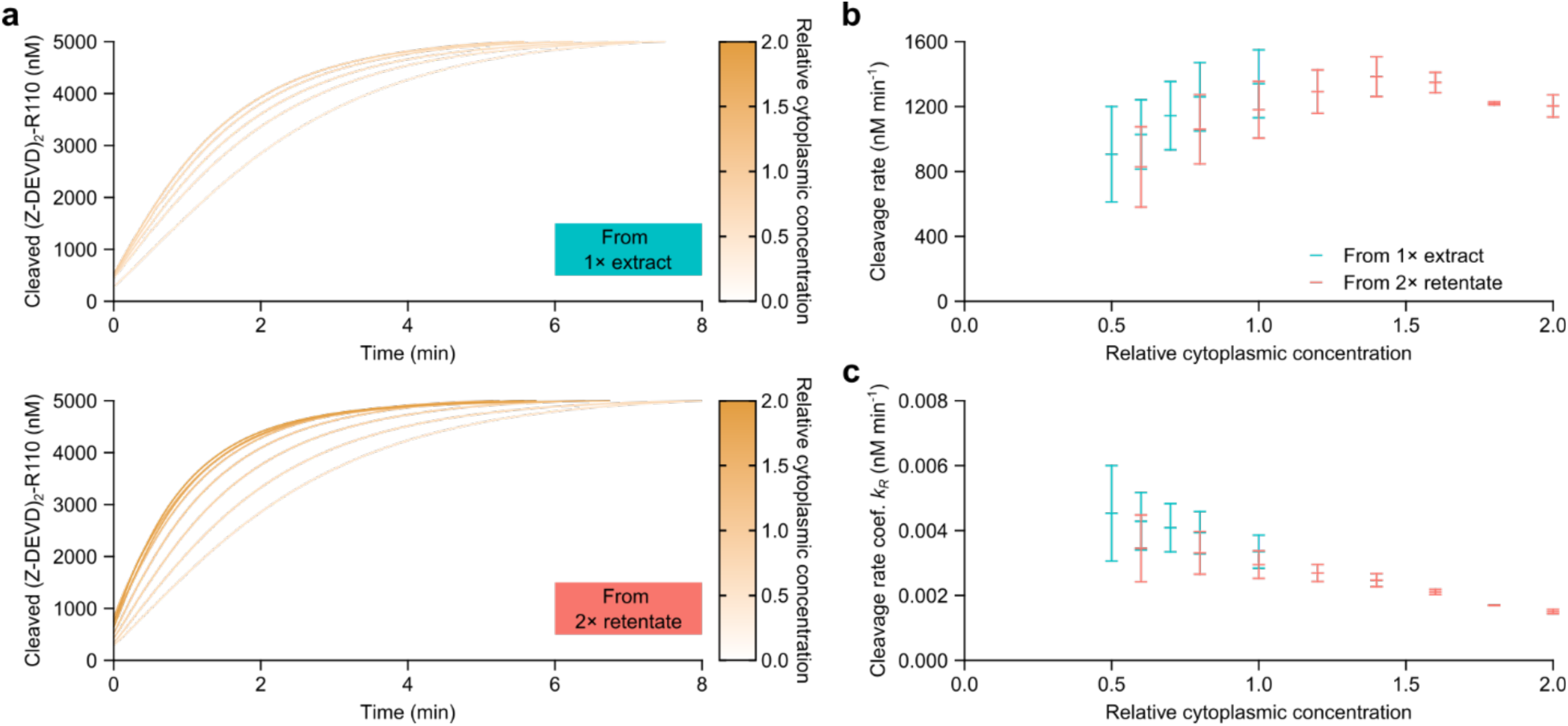
Measuring (Z-DEVD)_2_-R110 cleavage rate coefficient. 𝒌_𝑹_**. a** Kinetics of (Z-DEVD)_2_-R110 cleavage in freshly prepared apoptotic extracts at various cytoplasmic concentration. These extracts were prepared from 1x extract (top) or 2x retentate (bottom). **b** Cleavage rates of (Z-DEVD)_2_-R110 at different cytoplasmic concentrations were estimated based on the initial timepoints. **c** Second-order rate coefficients at different cytoplasmic concentrations were computed by accounting for (Z-DEVD)_2_-R110 concentration and nominal active caspase 3/7 concentrations. Means ± S.E.M. compiled from 3 independent samples are shown.

## Notes

### Competing Interest Statement

The authors have declared no competing interest.

